# CD11c+ CD8 T cells cause IFN-γ-dependent autoimmune neuroinflammation that is restrained by PD-1 signaling

**DOI:** 10.1101/2025.02.23.639743

**Authors:** Daniel Hwang, Larissa Lumi Watanabe Ishikawa, Maryam S. Seyedsadr, Soohwa Jang, Ezgi Kasimoglu, Abdolmohamad Rostami, Guang-Xian Zhang, Bogoljub Ciric

## Abstract

In multiple sclerosis (MS) lesions, CD8 T cells outnumber CD4 T cells, suggesting that they contribute to MS pathology. However, little is known on the role of CD8 T cells in MS, partially due to the nearly ubiquitous use of experimental autoimmune encephalomyelitis (EAE) models that are mediated by CD4 T cells, with no CD8 T cell contribution. Importantly, MS and EAE differ in both their distribution of CNS lesions and their symptoms, indicating differences in CNS inflammation. MS lesions are more prevalent in the brain, while EAE lesions are more frequent in the spinal cord. Additionally, neurologic deficits in MS rarely parallel the paralysis typical for CD4 T cell-mediated EAE (CD4-EAE). In contrast, CD8-EAE models suggest that CD8 T cells preferentially cause brain inflammation, but little is known regarding how brain and spinal cord inflammation may differ, and how CD8 T cells may contribute to those differences. We have established an adoptive CD8-EAE mouse model characterized by brain-centered inflammation, severe ataxia, and weight loss. CNS inflammation in the brain and spinal cord differed in immune cell numbers, cellular composition, and inflammatory signatures. CD8-EAE could be suppressed by blocking IFN-γ, and exacerbated by blocking PD-1, with concomitant changes in numbers of CNS-infiltrating monocytes. Most CD8 T cells in the CNS were CD11c^+^, suggesting that they are the pathogenic subset. We describe a robust CD8-EAE model, identify differences between brain and spinal cord inflammation, and characterize mechanisms that control CD8 T cell-mediated neuroinflammation, thereby furthering understanding of these cells in EAE/MS.

**Brief Summary:** CD8 T cells are understudied in MS due to a lack of suitable animal models. We developed a CD8 T cell dependent model of MS.

## INTRODUCTION

Multiple sclerosis (MS) is an autoimmune disease characterized by demyelinated lesions in the brain and, to a lesser extent, in the spinal cord [1]. Active MS lesions are laden with diverse immune cells, including monocytic cells, microglia, B cells, and both CD4 and CD8 T cells [2]. Amongst T cells in MS lesions, CD8 T cells are typically twice as numerous as CD4 T cells [2]. This is in stark contrast to conventional experimental autoimmune encephalomyelitis (EAE) models of MS, which all depend on CD4 T cells to initiate and sustain disease processes. CD8 T cells are neither present in EAE lesions, nor required for disease to fully develop [3]. The sole use of CD4 T cell-mediated EAE (CD4-EAE) models in developing MS therapeutics has resulted in a bias toward targeting CD4 T cell responses in MS. However, growing evidence suggests that CD8 T cells also contribute to MS pathology. Clinical trials in relapsing-remitting MS (RR-MS) targeting primarily CD4 T cell responses (anti-CD4 [4] and anti-IL-12/IL-23p40 [5] mAbs) failed to show clinical benefit, indicating that other cells, potentially CD8 T cells, are drivers of disease. There is also evidence of disease activity resulting from CD8 T cells; neuroantigen-specific, clonally-expanded CD8 T cells with highly differentiated phenotype are more frequent in MS patients than in healthy individuals [6]. Moreover, CD8 T cells in MS lesions express proinflammatory cytokines (e.g., IFN-γ, IL-17A, GM-CSF [7]) with similar frequency as CD4 T cells, further supporting possibility that CD8 T cells in MS play pathogenic role. Clinical and histological features of MS and EAE also differ in ways that could be attributed to pathogenic CD8 T cells. Most MS lesions are found in the brain, not the spinal cord [1]; causing an array of neurological deficits including impaired coordination and sensory defects [8]. This is reminiscent of findings by several seminal EAE studies showing that pathogenic myelin-specific CD8 T cells cause brain-centered inflammation, and impaired movement and coordination [9, 10]. This is decidedly different than in CD4-EAE models, in which lesions are predominantly in the spinal cord, with little involvement of the brain, and disease is characterized by ascending paralysis [11, 12]. These observations suggest that pathogenic mechanisms central to clinical manifestation of MS are not recapitulated in the CD4-EAE, despite their nearly ubiquitous use in MS research. Moreover, given that studies in CD8-EAE have found notable clinical similarities to MS strongly supports a view that CD8 T cells play an important pathogenic role in MS.

The dearth of knowledge on CD8 T cells in MS/EAE has primarily been caused by the lack of simple and reliable animal models in which CD8 T cells play a central role in causing robust CNS autoimmunity. Existing CD8-EAE models suffer from various experimental caveats, including reliance on CD4 T cells to induce disease [13, 14], or requiring radiation to render recipient mice susceptible to disease [10]. Other caveats include the transfer of pathogenic CD8 T cells directly into the CNS, thereby bypassing the need to cross the blood-brain-barrier (BBB) [10], or the use of vaccinia viruses to trigger CNS autoimmune disease [9]. Nevertheless, the insights gleaned from these studies have been highly valuable in revealing potential functions of CD8 T cells in MS. Myelin basic protein (MBP)-specific CD8 T cells from C3HeB/FeJ mice were shown to cause primarily brain lesions and a loss of coordinated movement that is referred to as atypical EAE [10]. Later, the T cell receptor (TCR) from a MBP_79-87_-specific CD8 T cell clone, named 8.8, due to expression of Vβ8 and Vα8 TCR, was used to generate TCR transgenic 8.8 mice [9]. These mice are largely tolerant for MBP_79-87_ [9], but can develop disease later in life [13]. The tolerance against MBP_79-87_ in 8.8 mice could be broken by viral infection with vaccinia virus, *via* a mechanism that involves spontaneous co-expression of TCRs specific for both vaccinia antigens and MBP_79-87_ on a subset of 8.8 CD8 T cells [9]. More recently, it was shown that 8.8 CD8 T cells could modify CD4-EAE by directing inflammation towards the brain, resulting in higher incidence of atypical EAE [14]. In these studies, however, CD4 T cells can obscure pathogenic mechanisms of CD8 T cells. For example, in early studies with MBP_79-87_-specific CD8 T cells, disease was IFN-γ-dependent, whereas IFN-γ was dispensable in CD4-EAE mice that received 8.8 CD8 T cells [14].

In the present study, we have developed a simple and reliable CD8-EAE model without irradiation of recipient mice, toxin injection, intrathecal transfer of CD8 T cells, viral infection, co-transfer of CD4 T cells, or transgenic pseudo autoantigens, which have been used in other CD8-EAE models. Our CD8-EAE model is based on transfer of *in vitro*-activated 8.8 T cells into naïve recipient mice, which then develop ataxia and weight loss, rather than ascending paralysis. This coincided with mostly brain inflammation, although the spinal cord was also affected. Interestingly, most of transferred 8.8 CD8 T cells become CD11c^+^, which has not been described previously in EAE or MS. Disease reproducibly started only several days after transfer of 8.8 CD8 T cells and was acute and monophasic, unless mice died at disease peak. The monophasic nature of disease was likely due to T cell exhaustion, as blocking PD-1 in recipient mice, or inducing memory phenotype in 8.8 CD8 T cells prior to transfer exacerbated disease. Lastly, transfer of 8.8 CD8 T cells caused influx of monocytes into the brain and spinal cord, which could be blocked by treatment with anti-IFN-γ mAb, thereby suppressing disease, suggesting that monocytes play an essential role in CD8-EAE pathology.

## MATERIALS AND METHODS

### Mice

8.8 mice (on C3HeB/Fe/J background) were generously provided by Dr. Joan Goverman. Wild type C3HeB/FeJ mice were obtained from Jackson Laboratories (Stock #000658).

### Induction of CD8-EAE

Splenocytes from 8.8 mice were isolated by mechanical dissociation followed by red blood cell lysis using RBC lysis buffer (Biolegend). Cells were plated at 2×10^6^ cells/mL in 6-well plates with 10 mL of IMDM + 2 mM L-glutamine and 10% heat-inactivated FBS. Cells were stimulated for 72 h with 0.33 μg/mL MBP_79-87_ and recombinant cytokines, as indicated in the results. Cells were washed twice then rested for 3 days in media with 5 ng/mL IL-2 (PeProTech). Each day cells were re-plated at 1×10^6^ cells/mL in fresh media supplemented with IL-2. For experiments where 2-deoxyglucose (2-DG) and rapamycin were used, 8.8 T cells were stimulated for 3 days in Tc1 conditions supplemented with 50 nM rapamycin or 4 mM 2-DG. During rest, 2-DG-treated cells were treated with 1 mM 2-DG.

In experiments in which cells were activated twice, cells were re-activated with anti-CD3 mAb (clone: 145-2C11; Bio X Cell) and anti-CD28 mAb (clone: 37.51; Bio X Cell). Plates were coated with anti-CD3 and anti-CD28 mAbs by covering the bottom of each well with 1 μg/mL anti-CD3 and 1 μg/mL anti-CD28 mAbs in PBS and incubating overnight at 37 ^o^C. Cultures were then restimulated for 2-3 days at 1×10^7^ cells/mL in 6-well plates with 10 mL media. CD8 T cells were harvested and purified by MACS separation by negative selection (Miltenyi Biotec) and resuspended in IMDM medium without FBS at 2×10^7^ cells/300 μL media that was supplemented with 0.2-0.4 μg IL-2 per mouse. 2×10^7^ cells were transferred by i.v. tail injection into recipient mice.

### Clinical Scoring

Mice were scored using atypical EAE scale as follows: grade 1, hindlimb clasping and/or hyperactivity; grade 2, hindlimb weakness and/or spastic movement; grade 3, spastic movement and dragging of hind limbs when walking (typically presenting as severe loss of coordinated movement, not paralysis); grade 4, head tilt with body leaning, forelimb weakness; grade 5, moribund or death.

### Cell Isolation and Flow Cytometry Analysis

To isolate the CNS mononuclear cells, blood was removed by cardiac perfusion with 60 mL PBS. The spinal cord was then flushed from the spinal column with PBS. The brains and spinal cords were combined and cut into small pieces, which were then incubated with Liberase TL (Roche) dissolved in RPMI at 0.7 mg/mL at 37 ^o^C for 30 min. The reaction was stopped by using complete media containing FBS, and the tissue was homogenized by pushing it through a 100 μm sterile filter with a syringe plunger. The homogenate was then centrifuged at 1500 RPM (300 x g) for 5 min and resuspended in 10 mL of 40% 1x Percoll-PBS (90% Percoll, 10% 10x PBS) and centrifuged at 2000 RPM without brake at room temperature for 30 min. The cells that pelleted were collected, diluted with PBS or media, and centrifuged at 1500 RPM (300 x g) for 5 min. The isolated cells were then stimulated with PMA (50 ng/mL; Sigma Aldrich), ionomycin (500 ng/mL; Sigma Aldrich), and 1 μL/mL Golgiplug (BD Biosciences) for 4 h at 37 ^o^C or stained without stimulation. After stimulation, the cells were washed with PBS containing 3% FBS (v/v). Cell surface antigens were then stained with antibodies in 100 μL of PBS/3% FBS for 20-30 min at 4 ^o^C. The cells were then washed and fixed with 100 μL of Fix and Perm Medium A (Thermo Fisher) for 20 min at room temperature and washed again. The cells were permeabilized with Fix and Perm Medium B (Thermo Fisher) and then stained with antibodies against intracellular antigens in 100 μL of Fix and Perm Medium B and 100 μL of PBS/3%FBS for 1 h or overnight. Finally, the cells were washed twice, resuspended in 500 μL PBS and analyzed on a BD FACSAria Fusion flow cytometer (BD Biosciences).

## RESULTS

### Adoptive transfer of 8.8 Tc1 cells causes severe EAE, whereas 8.8 Tc17 cells cause mild disease

In adoptive CD4-EAE, both Th1 and Th17 cells can cause neuroinflammatory disease [15]. In MS lesions, both CD4 and CD8 T cells produce IFN-γ and IL-17A [7], suggesting that both T cell subsets are pathogenic. Thus, we tested whether Tc1 and Tc17 cells could induce CNS autoimmunity. We polarized 8.8 CD8 T cells into either Tc1 or Tc17 lineage and expanded them to large numbers from small number of donor mice (Fig. 1A,B). Tc1 cells uniformly produced IFN-γ, while Tc17-polarized cells produced IL-17A, IFN-γ, or both cytokines. Adoptive transfer of 2×10^7^ Tc1 or Tc17 cells into naive recipients caused an acute and monophasic disease characterized by severe ataxia and rapid weight loss (Fig. 1C,D, Supplemental Video 1). Tc1 cells caused more severe clinical disease, weight loss, and CNS inflammation than Tc17 cells, (Fig. 1 C,D,E). Mice with CD8-EAE mediated by Tc1 cells had greater numbers of CD45^+^ cells in the brain *vs*. the spinal cord (Fig 1F). Unlike to ascending paralyses characteristic for CD4-EAE [1], mice that developed CD8-EAE retained the ability to move their tails and limbs, but lost coordinated movement to a varying degree, including loss of ability to right themselves. At disease peak, some mice became moribund or died (Supplemental Video 1), but those that survived always fully recovered. Interestingly, 40% of mice also developed severe swelling of the eyes (data not shown), with eye inflammation also following a monophasic disease course (Fig. 1G,H). Notably, Tc1 cells activated *in vitro* only once, followed by a rest and expansion period, had sufficient capacity to cause disease in recipient mice (Supplementary Fig. 1A,B). Similar to some reports about adoptive CD4-EAE [16], administration of pertussis toxin to recipient mice resulted in less severe CD8-EAE (Supplemental Fig. 1A,B). These data show that both Tc1- and Tc17-polarized 8.8 CD8 T cells can induce severe monophasic disease, starting only several days post transfer into naïve recipient mice. In contrast to Th1 and Th17 cells, Tc17 cells were notably less pathogenic than Tc1 cells, indicating that encephalitogenicity of CD8 T cells relies on IFN-γ, which is in direct contrast to suppressive role of IFN-γ in CD4-EAE [17]. Given that Tc1 cells were markedly more encephalitogenic than Tc17 cells, we continued our studies using Tc1-polarized 8.8 CD8 T cells.

**Figure 1.**
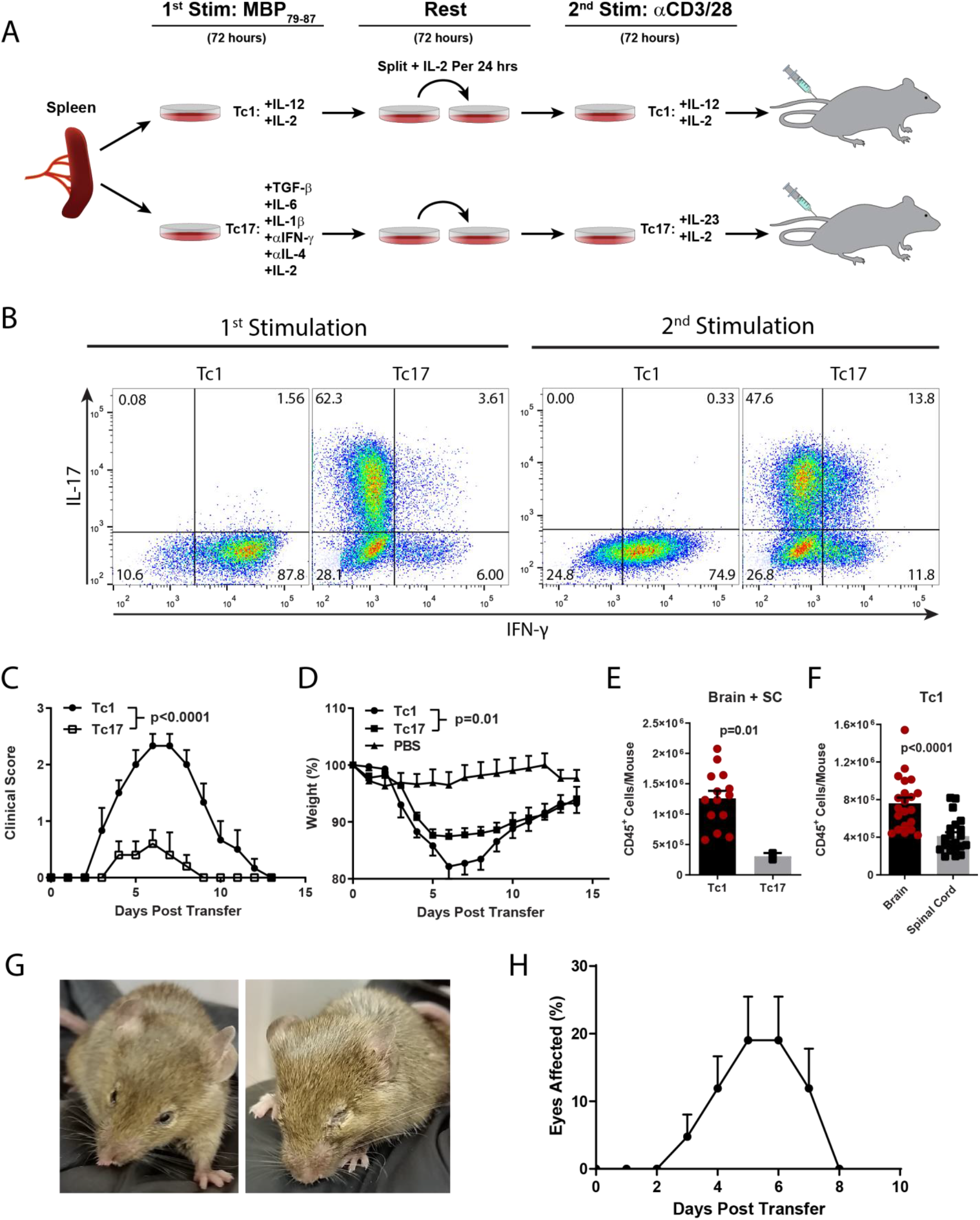
Adoptive transfer of 8.8 Tc1 and Tc17 cells induces autoimmune neuroinflammation. **A)** Splenocytes from 8.8 mice were stimulated with MBP_79-87_ for 3 days in either Tc1-, or Tc17-polarizing conditions. Tc17 condition were TGF-β (10 ng/mL), IL-6 (50 ng/mL), IL-1β (10 ng/mL), anti-IFN-γ (10 μg/mL), and anti-IL-4 (10 μg/mL). Tc1 condition were IL-12 (20 ng/mL) and IL-2 (5-10 ng/mL). After 72 h cells were rested in either 2 ng/mL IL-2 for Tc17 polarization, or 5-10 ng/mL IL-2 for Tc1 polarization. Each day cells were split and replated in fresh media supplemented with IL-2 at concentrations indicated above. After 3 days of rest, cells were reactivated for 2.5 days with plate-bound anti-CD3 and anti-CD28 in the presence of IL-23 (10 ng/mL) and IL-2 (2 ng/mL) for Tc17 cells, and IL-12 (2 ng/mL) and IL-2 (5-10 ng/mL) for Tc1 cells. For CD8-EAE induction, CD8^+^ cells were purified by negative selection using MACS beads and 2 x 10^7^ cells were transferred to recipient mice *via* tail vein injection. **B)** Flow cytometry plots showing Tc1- and Tc17-polarized 8.8 CD8^+^ T cells stained for IL-17A and IFN-γ after 1^st^ and 2^nd^ activation. **C)** 2 x 10^7^ MACS-purified CD8^+^ T cells and 0.2-0.4 μg recombinant murine IL-2 were i.v. transferred to naïve recipient mice. Clinical course for CD8-EAE (n=5-7 mice per group). Mice from two independent experiments are shown. **D)** Change in weight during CD8-EAE. Significance was determined by Two-way ANOVA. **E)** Number of CD45^+^ cells isolated from the CNS (brain + spinal cord). **F)** Number of CD45^+^ cells isolated from the brain or spinal cord. Significance was determined by t test. **G)** Photos of inflamed eyes during Tc1-mediated CD8-EAE. **H)** Average number of inflamed eyes over the course of CD8-EAE (n=21).

### Tc1 cells cause distinct inflammation in the brain and spinal cord

CD8 T cells have ben shown to mediate more severe inflammation in the brain than spinal cord [14], and our own data show that more immune cells are present in the brain during CD8-EAE (Fig. 1F). To characterize CNS inflammation mediated by 8.8 Tc1 cells, we have characterized the phenotype of immune cells from the brain and spinal cord at disease peak. CD45^+^ cells isolated from the CNS were virtually all CNS-resident immune cells and immune cells infiltrated into CNS tissue and were not contaminated with cells from the blood due to incomplete perfusion (Supplemental Fig. 2A). Among CD45^+^ cells from the CNS, the predominant cell types were CD8 T cells, monocytes and monocyte-derived cells, and microglia (Fig. 2A). CD8 T cells comprised the majority of lymphocytes in the CNS, with only 1-3% of immune cells being CD4^+^ T cells and CD19^+^ B cells (Fig. 2B). Over 95% of CD8^+^ cells were CD3^+^, and nearly all CD45^Lo^CD11b^+^ cells were Sall1^+^ microglia (Supplemental Fig. 3B-D). There were notable differences in immune cell composition and their phenotypes between the brain and spinal cord. MoDCs and MΦs, which are highly inflammatory myeloid cells in CD4-EAE models [18], were present in higher frequencies in the spinal cord compared to the brain. In the brain, the majority of myeloid cells were microglia (Fig. 2B). Myeloid cells in the spinal cord appeared to be more activated, with higher frequency of CD11c^+^ and MHC II^+^ monocytes, and nearly all microglia expressed CD11c (Fig. 2C,E). Notably, the majority of CD8 T cells also expressed CD11c (Fig. 2C). In both the brain and spinal cord, there were similar frequencies of monocytes expressing IFN-γ, TNF, and CCL2, but only few monocytes expressed IL-1β, CXCL9 and Arginase 1 (Fig. 2D). A higher frequency of microglia from the spinal cord expressed IFN-γ, with higher MFI for IFN-γ than in the brain (Fig. 2F-H). Monocytes from the brain and spinal cord had similar frequency of IFN-γ expression, but lower MFI for IFN-γ from the spinal cord (Fig. 2D, Supplemental Fig. 3E,F). Interestingly, a higher frequency of microglia expressed CCL2 in the spinal cord, suggesting that CCL2 secreted by microglia attracts more monocytes in the spinal cord (Fig. 2F). These data indicate that although overall infiltrating immune cell numbers are lower in the spinal cord, both brain and spinal cord inflammation are likely relevant to pathology in CD8-EAE.

**Figure 2.**
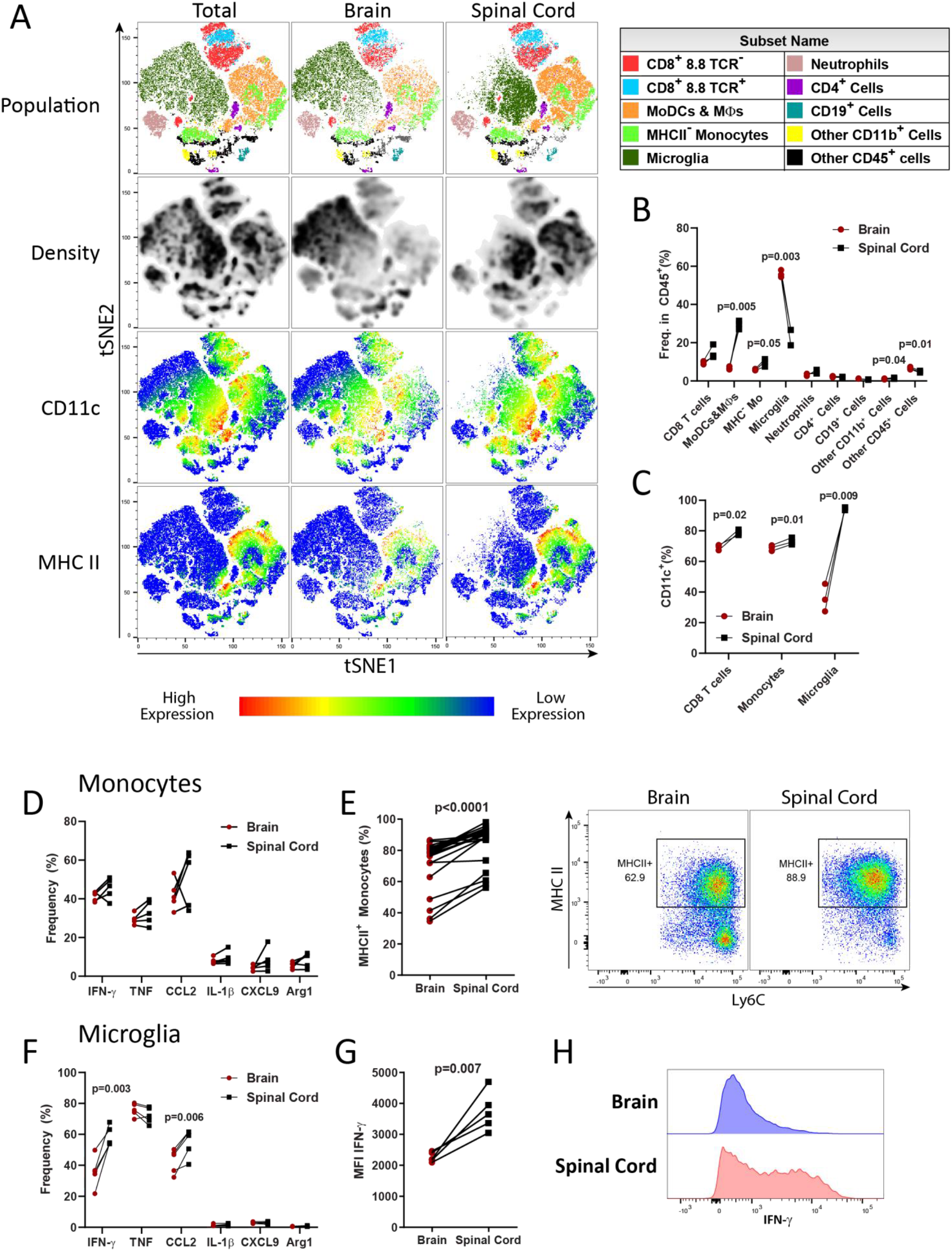
Tc1 cells cause distinct inflammation in the brain and spinal cord. **A)** Splenocytes from 8.8 mice were activated with MBP_79-87_ for 3 days in Tc1 condition, followed by 3 days expansion in IL-2 and CD8 T cell purification by MACS negative selection. 2 x 10^7^ cells were transferred i.v. into recipient mice. Mice were then sacrificed 7 days later and the brains and spinal cords analyzed separately by flow cytometry. Data from 1 (n=3) of 2 experiments with similar results are shown. CD45^+^ cells from the CNS of recipient mice were clustered by *t*SNE and populations are colored based on manual gating. CD8 T cells were defined as CD45^Hi^CD8^+^, with 8.8 TCR^+^ and 8.8 TCR^-^ cells being defined as TCR Vα8.3^+^Vβ8.1/8.2^+^ and TCR Vα8.3^-^Vβ8.1/8.2^-^, respectively. MoDCs and MΦs were defined as CD45^Hi^CD11b^+^Ly6G^Lo/-^Ly6C^+^MHCII^+^, microglia as CD45^Lo^CD11b^+^, neutrophils as CD45^+^Ly6G^Hi^CD11b^+^, CD4^+^ cells as CD45^Hi^CD11b^+^. CD19^+^ cells were CD45^Hi^CD19^+^, Other CD11b^+^ cells were CD45^Hi^CD11b^+^CD4^-^CD8^-^Ly6G^-^Ly6C^-^, Other CD45^+^ cells were CD45^+^CD4^-^CD8^-^CD11b^-^CD19^-^. **B)** Quantification of frequency of populations among CD45^+^ cells from A. **C)** Frequency of CD11c^+^ cells among CD45^Hi^CD8^+^ T cells, CD45^Hi^CD11b^+^Ly6G^-^Ly6C^+^ monocytes, and CD45^Lo^CD11b^+^ microglia. **D)** Expression of indicated cytokines and chemokines amongst monocytes from cells treated with PMA/Ionomycin/Golgiplug. **E)** Frequency of MHCII^+^ expression among monocytes. **F)** Expression of indicated cytokines and chemokines amongst microglia from cells treated with PMA/Ionomycin/Golgiplug. **G)** Median fluorescence intensity (MFI) of IFN-γ^+^ microglia. **H)** Histograms depicting IFN-γ expression in microglia. Significance was determined by paired t test.

### CD8 T cells in the CNS are predominantly CD11c^+^ and have inflammatory phenotype

We characterized the phenotype of CD8 T cells in the CNS during CD8-EAE. Approximately half of the CD8 T cells did not express transgenic 8.8 TCR (Fig. 3A). It is unclear whether these 8.8 TCR^-^ CD8 T cells were endogenous cells of recipient mice, or transferred 8.8 CD8 T cells that had downregulated expression of 8.8 TCR, but a portion of 8.8 TCR^-^ cells expressed either TCR Vβ8 or Vα8 chain, indicating that these cells derive from the host. Most CD8 T cells in the CNS had Tc1 phenotype, with over 80% of 8.8 TCR^+^ cells and approximately 70% of 8.8 TCR^-^ cells expressing IFN-γ (Fig. 3B). CD8 T cells also expressed GM-CSF, TNF, and CCL2, but 8.8 TCR^-^ cells had lower frequency of GM-CSF expression than 8.8 TCR^+^ cells. Unexpectedly, large proportions of CD8 T cells, both 8.8 TCR^+^ and 8.8 TCR^-^, expressed CD11c (Fig. 3C). Approximately 80% of 8.8 TCR^+^ cells were CD11c^+^, which is 6-fold enrichment in CD11c^+^ cells when compared to their frequency *in vitro*, prior to transfer into recipient mice. 8.8 TCR^-^ cells had lower frequency of expression and MFI for CD11c than 8.8 TCR^+^ cells (Fig. 3D,E). Consistent with reports that CD11c^+^ CD8 T cells have enhanced effector function, these cells had higher frequencies of IFN-γ and GM-CSF expression than CD11c^-^ cells (Fig. 3F). Our CD8-EAE model is characterized by monophasic disease, suggesting that inflammation is self-limiting. One possible explanation for this phenomenon is T cell exhaustion, which is commonly caused by PD-1 signaling. We characterized PD-1 and PD-L1 expression by immune cells from the CNS during CD8-EAE and found that PD-1 was abundantly expressed on all CD8 T cells, while myeloid cells expressed little PD-1 (Fig. 3G,H). Among CD45^+^ cells, MoDCs and MΦs expressed the highest levels of PD-L1, but CD8 T cells also uniformly expressed PD-L1, even more so than microglia, neutrophils and some monocytes. Interestingly, among CD8 T cells, 8.8 TCR^-^ cells had lower MFI for PD-1 and lower frequency of PD-1^+^ cells than 8.8 TCR^+^ cells (Fig. 3I, Supplemental Fig. 3A), indicating reduced exhaustion. Both 8.8 TCR^+^ and 8.8 TCR^-^ cells had similar MFI and frequency of PD-L1 expression (Supplemental Fig. 3B,C). Lastly, due to the increased frequency of MoDCs and MΦs in the spinal cord *vs*. the brain, there was increased MFI for PD-L1 among CD45^+^ cells in the spinal cord (Supplemental Fig. 3D). These data suggest that disease is self-limiting due to PD-1-mediated T cells exhaustion. Further, CD11c expression seems to characterize CD8 T cells with terminal effector function in the CNS during CD8-EAE.

**Figure 3.**
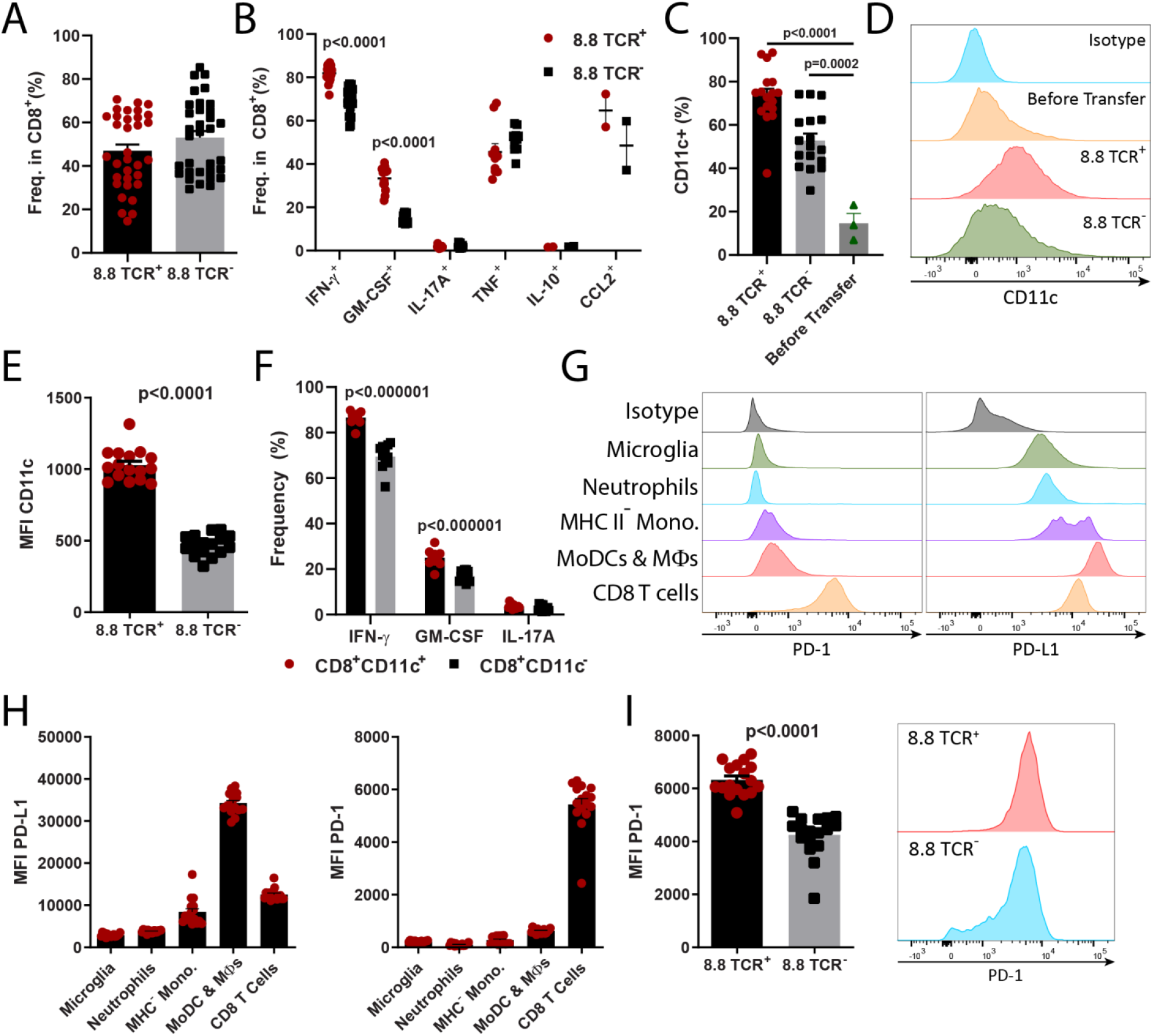
CD8 T Cells in the CNS during CD8-EAE are predominately CD11c^+^ with proinflammatory phenotype. CD8-EAE was induced by i.v. transfer of 2 x 10^7^ Tc1 cells and mice were sacrificed after 7 days. Brain and spinal cord cells were combined and analyzed by flow cytometry. **A)** Frequency of 8.8 TCR^+^ cells (TCR Vβ8.1/8.2^+^ TCR Vα8.3^+^) and 8.8 TCR^-^ cells among CD8^+^ cells. n=33 mice, from 5 experiments. **B)** Frequency of IFN-γ^+^, GM-CSF^+^, IL-17A^+^, TNF^+^, IL-10^+^, CCL2^+^ cells among 8.8 TCR^+^ CD8^+^ cells and 8.8 TCR^-^ CD8^+^ cells (n=2-22 mice) with data pooled from multiple experiments in all cases except for CCL2. **C)** Frequency of CD11c^+^ cells among 8.8 TCR^+^ CD8^+^ cells and 8.8 TCR^-^ CD8^+^ cells in the CNS (n=17 from 3 experiments) and *in vitro* before transfer (n=3 independent cultures). **D)** Histograms depicting expression of CD11c by CD8^+^ cells in the CNS and from *in vitro* cultures. **E)** MFI of CD11c on 8.8 TCR^+^ and 8.8 TCR^-^ CD8^+^ cells in the CNS (n=16 from 2 experiments). **F)** Frequency of IFN-γ^+^, GM-CSF^+^, and IL-17A^+^ cells among CD11c^+^CD8^+^ and CD11c^-^CD8^+^ cells (n=12 mice). **G)** Histograms depicting expression of PD-1 and PD-L1 by microglia, neutrophils, MHC^-^ monocytes, MoDCs and MΦs, and CD8 T cells. **H)** MFI of PD-L1 and PD-1 among indicated cell populations. **I)** MFI for PD-1 among 8.8 TCR^+^ and 8.8 TCR^-^ CD8^+^ T cells and histograms depicting PD-1 expression. For H-I, n=16 mice from 2 experiments. Significance was determined by unpaired t test.

### Inflammation during CD8-EAE is restrained by T cell exhaustion

High expression of PD-1 on CD8 T cells in the CNS suggested that neuroinflammation in CD8-EAE is curtailed by CD8 T cell exhaustion. To test this hypothesis, we blocked PD-1 signaling during EAE using neutralizing mAbs against either PD-1 or PD-L1. Blocking PD-1 signaling with mAbs against PD-1 or PD-L1 exacerbated motor dysfunction and eye inflammation during CD8-EAE (Fig. 4A,B; Supplemental Fig. 4A,B). Blocking PD-1 increased numbers of immune cells in the brain, and to a greater extent in the spinal cord (Fig. 4C) Characterization of immune cells from the brain and spinal cord showed that the increase in cell numbers was primarily caused by an increase in inflammatory monocyte-derived cells (Fig. 4D E). αPD-1 treatment reduced the MFI of PD-1 on CD8 T cells in the CNS, indicating reduced T cell exhaustion (Fig. 4F,G). αPD-1 treatment did not affect PD-L1 expression on any CD45^+^ cells in the CNS (Supplemental Fig. 4C,D). Lastly, we found that exacerbated disease in αPD-1-treated mice correlated with an increase in 8.8 TCR^-^ CD8 T cells (Fig. 4H). Together, these data indicate that disease severity in this EAE model is limited by CD8 T cell exhaustion.

**Figure 4.**
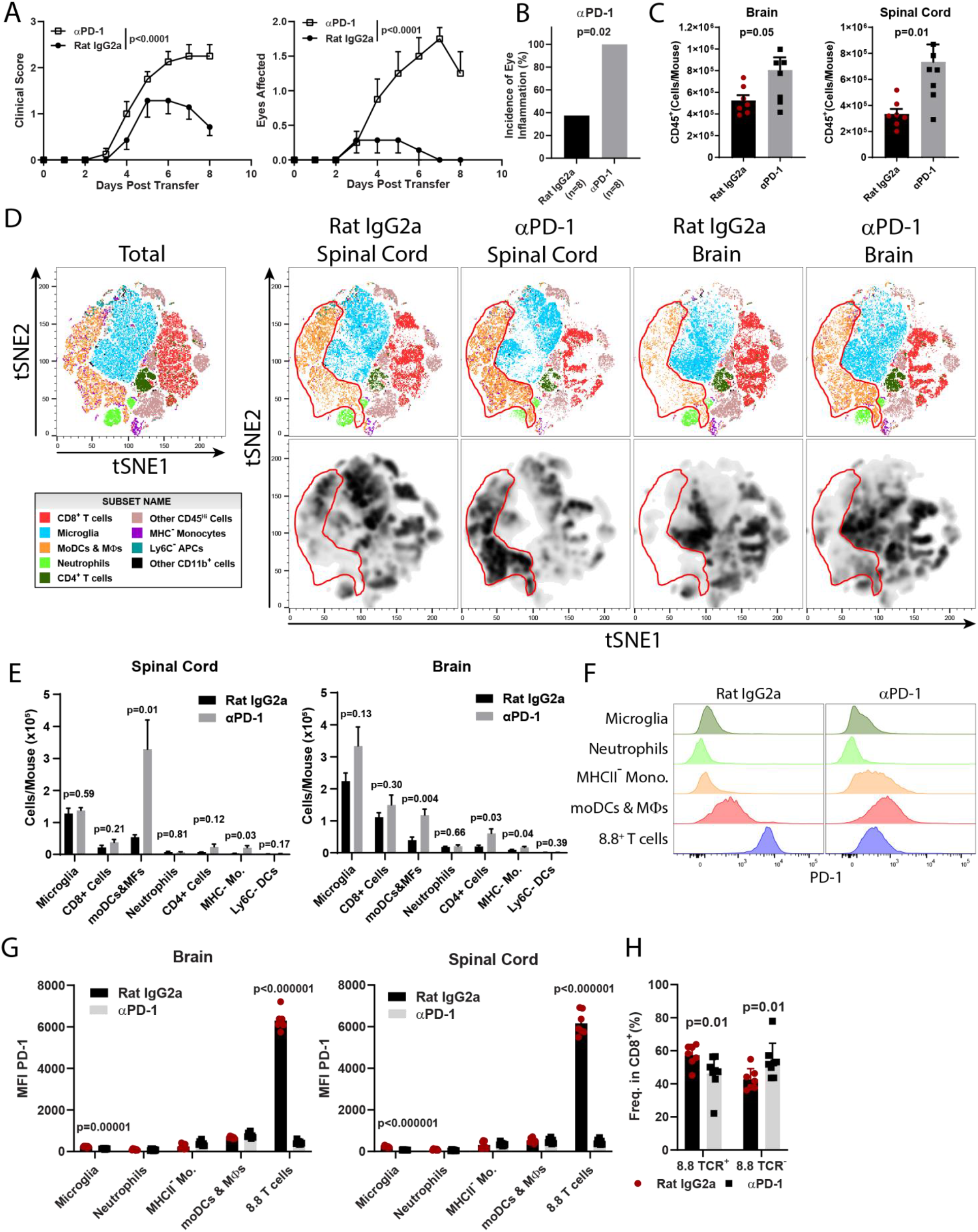
Blocking PD-1 signaling exacerbates CD8-EAE. CD8-EAE was induced, and recipient mice were treated with 200 μg αPD-1 or isotype mAb (Rat IgG2a) once per day starting from day of CD8-EAE induction (n=7-8 mice per group; compiled from 3 independent experiments). **A)** Clinical scores and number of inflamed eyes over time are shown. Statistical significance was determined by two-way ANOVA. **B)** Incidence of eye inflammation in mice treated with αPD-1 or isotype mAb. Significance was determined by Chi-squared test. **C)** Number of CD45^+^ cells from the brain and spinal cord of mice treated with αPD-1 and sacrificed on day 8 after CD8-EAE induction. **D)** tSNE clustering of CD45^+^ cells from high color flow cytometry data. CD8^+^ T cells were defined as CD8^+^CD3^+^; microglia as CD11b^+^Sall1^+^; MoDCs and macrophagess as Sall1^-^CD11b^+^Ly6G^lo/-^ Ly6C^+^MHCII^+^; neutrophils as Sall1^-^CD11b^+^Ly6G^Hi^Ly6C^Int^; CD4 T cells as CD4^+^CD3^+^. Other CD45^Hi^ cells did not express lineage markers that were stained in this panel. MHC^-^ monocytes were Sall1^-^CD11b^+^Ly6G^lo/-^ Ly6C^+^MHCII^-^;Ly6C^-^ APCs were Sall1^-^CD11b^+^Ly6C^-^MHCII^+^; other CD11b^+^ cells did not express Sall1, Ly6C, Ly6G, and MHCII. **E)** Quantification of cell numbers based on gating from D. **F)** Histograms depicting expression of PD-1 among cell populations. **G)** MFI for PD-1 in the brain and spinal cord among populations from F. **H)** Frequency of 8.8 TCR^+^ and 8.8 TCR^-^ cells among CD8^+^ cells. For C-H statistical significance was determined by two-tailed unpaired t test.

We noted some variability in disease severity among CD8-EAE experiments, including the development of fulminant disease that resulted in rapid death of recipient animals in some experiments. Over the course of our investigation, we noted that some 8.8 mice spontaneously develop overt symptoms of EAE, which agrees with a recently published report [13]. The occurrence of spontaneous EAE (sEAE) demonstrates a break in immune tolerance in some 8.8 mice that leads to the development of pathogenic CD8 T cells. We therefore hypothesized that the variability in disease severity among repeat experiments was due to differences in phenotype of CD8 T cells harvested from donor 8.8 mice for adoptive transfer. In this view, CD8 T cells from mice that developed sEAE would have the most pathogenic phenotype. Although animals that developed sEAE were rare, we have included *in vitro* analyses of a mouse with sEAE. We activated splenocytes from this mouse *in vitro* and compared their phenotype to cells from mice that appeared healthy. Cells from the sEAE mouse had more activated and less exhausted phenotype, with decreased expression of PD-1, and increased expression of IL-2 and TNF. TIM-3 and IFN-γ expression was similar between sEAE mouse and the normal mouse (Supplemental Fig. 4E). Although these observations need to be validated by comparing cells from larger numbers of healthy mice with those that have developed sEAE, our data thus far suggest that immune status of naïve donor mice is not uniform and could cause variability in CD8-EAE.

### Rapamycin-treated 8.8 CD8 T cells have enhanced encephalitogenicity

Our data on the phenotype of CD8 T cells and blocking PD-1/PD-L1 suggest that the monophasic disease course in our CD8-EAE model is a consequence of CD8 T cell exhaustion that leads to elimination/anergy of CD8 T cells and resolution of CNS inflammation. Terminal exhaustion of effector CD8 T cells and an extension of immune responses mediated by them can be achieved by switching their phenotype from effector to memory [19]. We tested whether induction of memory-like cells could exacerbate or extend disease in our CD8-EAE model by treating 8.8 CD8 T cells during *in vitro* activation with rapamycin or 2-DG, which have been shown to induce a memory phenotype in CD8 T cells with enhanced and prolonged effector function *in vivo* [20, 21]. Rapamycin and 2-DG treatments differed in their ability to modify phenotype of 8.8 CD8 T cells. Rapamycin strongly enhanced production of inflammatory cytokines and chemokines, including IFN-γ, TNF, GM-CSF and CCL2, whereas 2-DG only enhanced GM-CSF production (Fig. 5A-E). Enhanced production of IL-2 by CD8 T cells is associated with memory phenotype and better survival [22]. Both rapamycin and 2-DG treatments increased frequency of IL-2-expressing CD8 T cells during their culture (Fig. 5F,G). The treatments also affected expression of exhaustion markers PD-1 and TIM-3, with rapamycin reducing expression of both PD-1 and TIM-3, whereas 2-DG did not affect PD-1 expression but markedly reduced frequency of TIM-3-expressing CD8 T cells (Fig. 5H,I). Treatment with 2-DG, but not rapamycin, upregulated TCF-1, a marker of stem cell memory T cells (Supplemental Fig. 5A). We then transferred 2-DG- and rapamycin-treated 8.8 CD8 T cells to recipient mice to induce CD8-EAE. Rapamycin-treated cells induced exacerbated disease when compared to control Tc1 cells, whereas 2-DG-treated cells were not encephalitogenic (Fig. 5J, Supplemental Fig. 5B). Even though manipulation of CD8 T cell phenotype in the context of CD8-EAE requires further exploration, our data nonetheless suggest that reducing exhaustion of CD8 T cells by inducing memory-like phenotype could enhance their pathogenicity.

**Figure 5.**
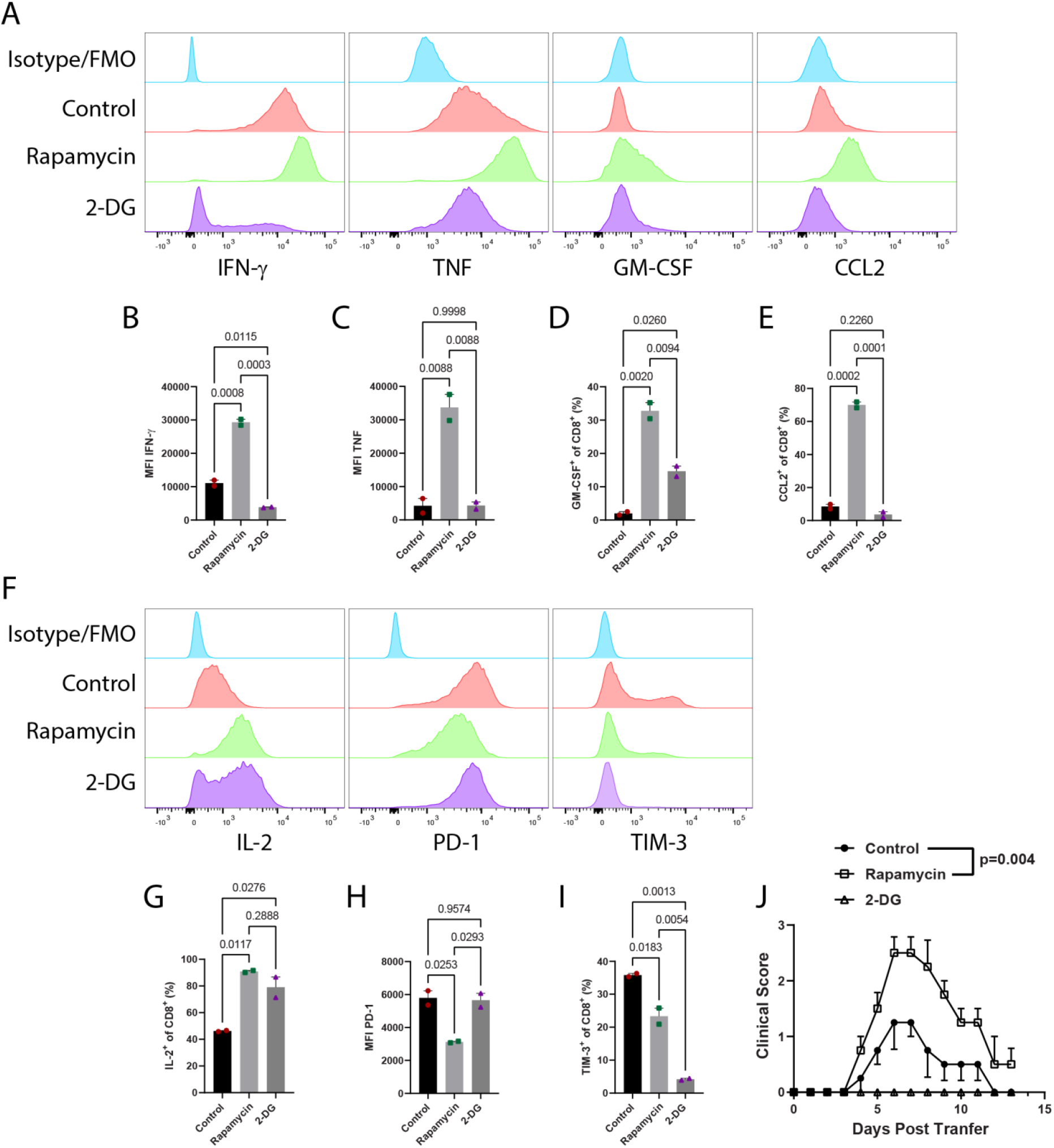
Rapamycin-treated 8.8 CD8 T cells induce more severe disease. 8.8 CD8 T cells were activated in Tc1 conditions with 50 nM rapamycin (Rapa) or 4 mM 2-deoxyglucose (2-DG). **A)** Expression of IFN-γ, TNF, GM-CSF, CCL2 in CD8^+^ cells after 3-day stimulation with MBP_79-87_. **B-C)** MFI of IFN-γ and TNF among IFN-γ^+^ and TNF^+^ cells. **D-E)** Frequency of GM-CSF^+^ and CCL2^+^ cells among CD8^+^ cells. **F)** Expression of IL-2, TIM-3 and PD-1 by CD8^+^ cells. **G)** MFI for PD-1 on CD8^+^ cells. **H)** Frequency of IL-2^+^ cells among CD8^+^ cells. **I)** Frequency of TIM-3^+^ cells among CD8^+^ cells. Significance for B-E, G-I was determined by one-way ANOVA with multiple comparisons testing. **J)** Following 1^st^ activation, cells were rested/expanded for 3 days in IL-2 media supplemented with 50 nM rapamycin or 1 mM 2-DG. 2 x 10^7^ CD8 T cells were then transferred to recipient mice to induce CD8-EAE. Clinical course is shown; n=4 for control and rapamycin; n=2 for 2-DG. Significance for J was determined by two-way repeated measures ANOVA.

### CD8-EAE pathology is dependent on IFN-γ

We next sought to determine which mechanisms control pathology in CD8-EAE. GM-CSF is essential for CD4-EAE development [23], but its role in CD8-EAE is unknown. Blocking GM-CSF with mAb moderately suppressed disease (Supplemental Fig. 6A), indicating that GM-CSF contributes to pathology in CD8-EAE, but, unlike in CD4-EAE, its role is not essential. It has been shown that Fas/FasL interactions are required for CD8 T cells to augment brain inflammation in an adoptive transfer model of CD4-EAE [14]. We tested whether Fas/FasL interactions play a role in CD8-EAE, but blocking FasL with mAb had no effect on disease course (Supplemental Fig 6B). In earlier studies, intrathecal transfer of MBP-specific CD8 T cells caused neuroinflammatory disease that could be blocked by anti-IFN-γ mAb [10]. We tested whether IFN-γ is also important in our CD8-EAE model and found that disease was potently suppressed by i.p. injections of anti-IFN-γ mAb (Fig. 6A). Mice receiving anti-IFN-γ mAb developed markedly less severe disease with less weight loss than animals treated with control IgG. Anti-IFN-γ mAb even blocked fulminant disease in experiments in which nearly all control animals quickly succumbed to CD8-EAE. In agreement with clinical disease suppression, mice treated with anti-IFN-γ mAb had reduced numbers of CD45^+^ cells in both the brain and spinal cord (Fig. 6B). We then sought to characterize the effects that blocking IFN-γ had on CNS inflammation. Clustering of CD45^+^ cells from the brain and spinal cords revealed reduced frequencies of moDCs and MΦs in both the brain and spinal cord (Fig. 6C,D). CD11c-expressing microglia in both the brain and spinal cord were also reduced (Fig. 6E), suggesting diminished microglia activation in anti-IFN-γ mAb-treated CD8-EAE mice. CD8 T cells were also reduced in frequency in the brain, but not in the spinal cord. CD8 T cells from anti-IFN-γ mAb-treated animals had reduced expression of PD-1, indicating reduced activation (Fig. 6F). Expression of PD-L1 among both CD11b^+^ cells and CD45^+^ cells was reduced, owing primarily to fewer moDCs and MΦs in the CNS (Fig. 6G, Supplemental Fig. 6). These data indicate that IFN-γ promotes infiltration of monocytes in the CNS and further suggests that monocyte are a major effector cell type of neuroinflammation in CD8-EAE.

**Figure 6.**
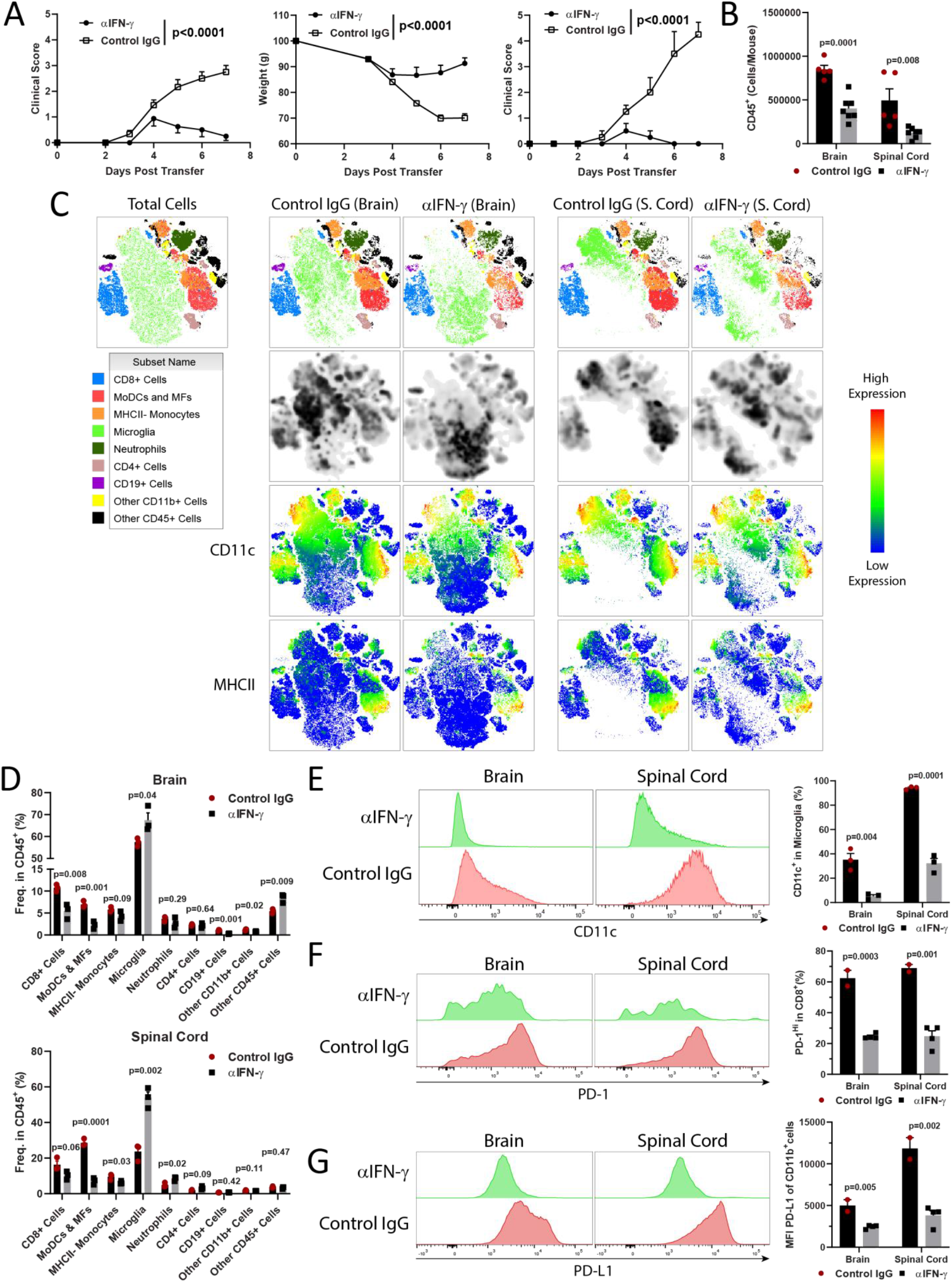
Blocking IFN-γ suppresses CD8-EAE. CD8-EAE was induced by transfer of 2 x 10^7^ CD8 T cells cultured in Tc1 conditions for 3 days then rested/expanded for 3 days before transfer. **A)** Clinical scores and change in weight for CD8-EAE mice treated with 200 μg/day anti-IFN-γ mAb or control IgG; n=8 per group, with data compiled from 2 experiments. Right panel shows clinical scores for a second independent experiment where CD8-EAE mice developed fulminant disease and were treated with either anti-IFN-γ or control IgG. n=4 per group. Significance was determined by two-way ANOVA. **B)** Numbers of CD45^+^ cells isolated from the brain and spinal cord on day 7 post transfer; n=5-8 per group compiled from 2 experiments. **C)** *t*SNE clustering of CD45^+^ cells isolated from the brain and spinal cord. CD8^+^ T cells were defined as CD45^+^CD8^+^; MoDCs and MFs as D45^Hi^CD11b^+^Ly6G^lo/-^Ly6C^+^MHCII^+^, neutrophils as CD45^Hi^CD11b^+^Ly6G^Hi^Ly6C^Int^; MHC^-^ monocytes as CD45^Hi^CD11b^+^Ly6G^lo/-^Ly6C^+^MHCII^-^; microglia as CD45^Lo^CD11b^+^; CD4 T cells as CD45^Hi^CD4^++^. CD19^+^ cells were defined as CD45^+^CD19^+^CD4^-^CD8^-^. Other CD11b^+^ cells were defined as CD45^+^CD11b^+^ cells that did not express other markers. Other CD45^+^ cells did not express other lineage markers that were stained in this panel. **D)** Histograms depicting expression of CD11c on microglia in the brain and spinal cord. **E)** Expression of PD-1 and PD-L1 on total CD45^+^ cells and on CD8^+^ cells from the CNS of mice treated with anti-IFN-γ or control IgG. **F)** Expression of PD-L1 on CD11b^+^ cells. Significance for C-F was determined by unpaired t test.

## DISCUSSION

We have developed an adoptive CD8-EAE model that relies on transfer of CD8 T cells that express transgenic TCR specific for MBP. The model is simple, requiring a single activation of the CD8 T cells *in vitro* before their i.v. transfer into naïve wild-type recipient mice, without even *P*. toxin injections. This is uncommonly minimalistic approach for a CD8-EAE model, as those described in the literature require additional manipulations to induce CNS autoimmunity, such as: irradiation of recipient mice, transgenic recipient mice that express pseudo-autoantigens, injection of CD8 T cells directly into the CNS, viral infection, etc. The drawback of these more complex models is not only added technical difficulty, but, also, that they are further removed from physiological processes/mechanisms, therefore representing them less reliably than our model. Further, our model is robust, as mice reliably develop clinical disease only four days after receiving CD8 T cells, disease peaking seven days after T cell transfer, and mice fully recovering two weeks after disease induction. In our experience, after optimization of multiple parameters, we have devised a simple protocol that reproducibly induces disease of moderate/high severity, without many mouse deaths due to overly severe disease, as was the case in our early experiments. Like in other CD8-EAE models, mice develop varied outward signs of disease, which could cause difficulty in scoring disease severity. However, a measurable parameter that objectively reflects disease severity was loss of body weight. Reassuringly, there was a good correlation of clinical score and loss of body weight, validating scoring scale as a realistic estimation of disease severity. As regards to the usefulness of our CD8-EAE model, we believe its simplicity, reproducibility, and physiological nature could be very useful in studying mechanisms of CD8 T cell-mediated neuroinflammation, as some of our experiments shown here demonstrate (e.g., role of IFN-γ). From the perspective of studying potential therapies that target CD8 T cell-mediated pathogenic processes, due to self-resolving disease course, therapeutic studies would be limited to prophylactic treatment regimen, serving as a predictor for therapeutic effects when treatments were started during ongoing disease. This limitation could hopefully be overcome by devising an approach to extend disease, as we attempted by treating 8.8 CD8 T cells with rapamycin and 2-DG. Still, our CD8-EAE model could be combined with a CD4-EAE model to generate hybrid CD8/CD4-EAE model that would perhaps represent MS better than individual CD8- and CD4-EAE models. Such hybrid EAE model has been explored, using transfer of naïve 8.8 CD8 T cells into mice with CD4 T cell-initiated EAE [14]. In this scenario, the development of autoreactive CD8 T cell response is secondary to CD4 T cell-induced neuroinflammation, with comparatively few CD8 T cells in the CNS when compared to CD4 T cells. However, this might not be the case in MS, and co-transfer of activated 8.8 CD8 T cells together with encephalitogenic CD4 T cells could represent MS pathogenesis more closely. Owing to persistent neuroinflammation driven by CD4 T cells, such a hybrid EAE model would not be monophasic, making it useful for testing therapies in the context of CNS autoimmunity driven by both CD4 and CD8 T cells, which likely occurs in most MS cases.

Experimental findings suggest that myelin-specific CD8 T cells direct CNS inflammation toward the brain, as transfer of autoreactive CD8 T cells in CD4-EAE increased brain inflammation, resulting in increased incidence of atypical EAE (loss of coordinated movement) [14]. It is unknown why CD8 T cells cause more brain inflammation than CD4 T cells, but it may involve differential expression of adhesion molecules. Melanoma cell adhesion molecule (MCAM) is expressed by a subset of human effector CD8 T cells, and blocking its activity reduces transmigration in a human BBB model [24]. Our own data supports the concept that certain subsets of CD8 T cells express adhesion molecules that could bias their accumulation in the brain. In the CNS of CD8-EAE mice, ∼80% of CD8 T cells expressed CD11c,far exceeding their systemic frequency (∼5%; [25]) and their frequency prior to transfer (∼15%) into recipient mice. CD11c^+^ CD8 T cells have not been described previously in the context of CNS autoimmunity, neither in EAE nor in MS. It would be highly relevant to determine whether CD8 T cells in some other CD8-EAE models also express CD11c, and, particularly, if CD8 T cells in MS lesions are CD11c+. At this time it is unclear whether CD11c upregulation on CD8 T cells is peculiarity of our model, or common occurrence among CD8 T cells in inflamed CNS areas. It is also unknown if and in what way CD11c might be important for CD8 T cell functions in CNS autoimmunity. CD11c, also known as α_X_ integrin, pairs with β_2_ integrin to form complement receptor 4 (CR4). In addition to myeloid dendritic cells, CR4 can be expressed on lymphocytes, including T cells [26, 27]. Knockout of CD11c (and therefore CR4) on T cells suppressed adoptive CD4-EAE [28]. Thus, it seems possible that CD11c directly contributes to the pathogenicity of autoreactive CD8 T cells in EAE. However, it is unlikely that CD11c+ and CD11c-CD8 T cells only differ in the expression of this molecule. It has been shown in viral infection that CD11c^+^ CD8 T cells have greater inflammatory effector function than their CD11c^-^ counterparts [29]. The effector functions of CD11c^+^CD8^+^ T cells appears to be centered on IFN-γ production, rather than on their cytolytic function [25], which is in agreement with our data showing that CD8-EAE is IFN-γ-dependent. Characterization of phenotypic and functional differences between CD11c+ and CD11c-CD8 T cells would be relevant not only in the context CNS autoimmunity, given that little is known on this topic in general. It appears that our CD8-EAE model could be useful in such studies.

In CD4-EAE, both Th1 and Th17 cells are sufficiently pathogenic to induce EAE [30]. Th1 and Th17 cells are present in MS lesions, which together with observations from EAE, indicate that both Th subsets are pathogenic in MS. Similarly, Tc1 and Tc17 cells can be found in MS lesions [7], but it is unknown whether these CD8 T cell subsets differ in their pathogenicity, because there are no studies directly testing encephalitogenicity of Tc17 cells. It was described, in an atypical active EAE model, that Tc17 cells themselves do not induce EAE, but provide crucial help in differentiation of encephalitogenic Th17 cells, which then cause the development of EAE [31]. It is unclear if, and to what extent, this indirect pro-encephalitogenic function of Tc17 cells are relevant in other EAE models.

In our testing of Tc1 *vs*. Tc17 cells in CD8-EAE, we were surprised to find that Tc1 cells were notably more encephalitogenic than Tc17 cells. Although we do not know why this may be, a plausible explanation could be that more abundant IFN-γ production by Tc1 cells renders them more pathogenic. Indeed, we found that IFN-γ was an essential pathogenic mediator in CD8-EAE, as neutralization of IFN-γ abrogated even fulminant disease. This agrees with other studies showing that CD8-EAE development is dependent on IFN-γ [10, 32, 33]. However, the ability for 8.8 CD8 T cells to promote brain inflammation in a model of CD4-EAE was independent on IFN-γ [14]. This is opposite to the protective role of IFN-γ in CD4-EAE, with IFN-γ knockout mice developing exacerbated disease [34]. In contrast, IFN-γ appears to be proinflammatory in MS, as administration of IFN-γ to MS patients during a clinical trial in the 1980’s caused an increase in MS relapse rate [35]. Thus, the role of IFN-γ in MS is likely better recapitulated in CD8-EAE than in CD4-EAE; implying that a relevant pathogenic mechanism of CD8 T cells in MS is their production of IFN-γ.

It is unknown how IFN-γ facilitates CD8-EAE, but our data indicate that IFN-γ participates in monocyte activation and recruitment into the CNS,As blocking IFN-γ greatly reduced numbers of monocytic cells in both brain and spinal cord. One potentially relevant mechanism is IFN-γ-dependent production of CCL2, which can be produced both by non-immune cells, including astrocytes [36-39], and immune cells, including monocytes and microglia [40, 41]. In agreement with this hypothesis is that we found widespread CCL2 production by CNS immune cells in CD8-EAE, including monocytes, microglia and CD8 T cells. Since blocking IFN-γ reduced numbers of inflammatory monocytes, and they are a source of CCL2, this may have had reduced monocyte recruitment to the CNS. Thus, it is possible that CD8 T cell-dervied IFN-γ is involved in inflammatory myeloid cell responses in the CNS during CD8-EAE.

It is not known if cytotoxicity of Tc cells contributes to their encephalitogenicity, but if it is important, then low encephalitogenicity of Tc17 cells could also be explained by lack of cytotoxicity. We and others have shown that Tc17 cells do not express granzyme B, perforin and are not cytotoxic [31, 42, 43], although perforin-independent [43] and FasL-dependent [44] cytotoxicity of Tc17 cells has been described. Perforin produced by CD8 T cells is important in disruption of BBB and development of brain inflammation [45]. Pharmacological inhibition of perforin largely reversed neurologic deficit caused by autoreactive CD8 T cells that caused BBB disruption [46]. Information on the role of granzyme B in EAE, and especially CD8-EAE is scant. It has been shown that inhibition of granzyme B in CD4-EAE reduced axonal and neuronal injury and maintained myelin integrity but did not reduce infiltration of CD4 and CD8 T cells into the CNS [47]. CD8 T cells expressing Granzyme B were observed in proximity or attached to oligodendrocytes or demyelinated axons, indicating that granzyme B contributes to their pathogenic function [48]. A recent study has shown that MS patients with progressive disease course have greater numbers of CD8 T cells with granzyme B expression than patients with relapsing remitting MS, and its expression positively correlated with disability and progression of MS [49]. T-bet was required for granzyme B expression in these CD8+ cells [49]. Overall, direct evidence for the role of cytotoxic molecules in encephalitogenicity of CD8 T cells is lacking, but it seems possible that they could be essential for EAE to develop when encephalitogenic CD4 T cells are absent and CD8 T cells alone need to initiate and drive CNS inflammation. Even though Th17 cells are highly encephalitogenic, this function is dependent on upregulation of T-bet expression in Th17 cells and switch of their phenotype toward Th1 [50]. Tc17 cells do not express T-bet, but they can upregulate its expression and switch to Tc1-like, ex-Tc17, phenotype [42], analogous to Th17/ex-Th17 cells. We have also seen in our experiments with 8.8 Tc17 cells that after 2^nd^ stimulation a portion of them co-expresses IL-17 and IFN-g, which signifies ongoing Tc17 to Tc1 switch. It is likely that this switching continues or even accelerates after transfer of Tc17 cells into recipient mice. However, in contrast to Th17 cells, this switch does not seem to impart encephalitogenicity to Tc17/ex-Tc17 cells. Our findings indicate that further study on encephalitogenicity of myelin-specific Tc17 cells is needed to answer questions that have arisen from our data.

Disease in our CD8-EAE model was monophasic, which correlated with high degree of PD-1 expression on CD8 T cells both *in vivo* and *in vitro*, indicating that T cell exhaustion contributes to self-limiting disease course. Accordingly, blockade of PD-1 or its ligand PD-L1 exacerbated ataxia in CD8-EAE. It has been shown that T cell exhaustion also limits CD4-EAE severity, as mice deficient for PD-1 or PD-L1 (but not PD-L2) develop more severe disease [51]. Our CD8-EAE disease was exacerbated by transfer of rapamycin-induced memory CD8 T cells; transferred cells expressed less PD-1 and more inflammatory cytokines, indicating that they are less exhausted. Thus, 8.8 T cell exhaustion is likely the primary reason for the monophasic nature off this EAE model, although we cannot exclude possibility that other immunoregulatory mechanisms also contribute to the resolution of neuroinflammation The monophasic disease course could be explained by the absence of Th responses in this model, which are required for sustained CD8 T cell responses across various diseases and therapeutic settings [52]. Our understanding of the CD8 T cell role in MS may therefore benefit from comparison between CD8-EAE and mixed CD4/CD8-EAE models, which could help delineate how CD4 T cells impact disease with a notable CD8 T cell component.

In conclusion, we have established a convenient adoptive CD8-EAE model, characterized by inflammation in both the brain and spinal cord. The simplicity of the model, namely use of naïve wild-type recipient mice, is unique among CD8-EAE models and more closely mimics physiological processes than other models. This could provide more realistic and therefore more relevant insights into processes occurring in MS. The neurological deficits in our model are different, and perhaps more similar to MS, than in common CD4-EAE models. This is likely caused by more severe brain inflammation than in CD4-EAE models, although specific nature of CD8 T cell-mediated inflammation, including its self-resolving course, could also have an impact on observable clinical deficits. We believe that our CD8-EAE model is a highly useful tool for studying an aspect of CNS autoimmunity that has been understudied, even though it is decidedly relevant for MS. Hence, using our model to study neuroinflammation could lead to better understanding of MS and open new avenues toward its therapy.

## Acknowledgements

We thank Joan Goverman for the kind gift of 8.8 mice.

## Funding

This work was supported by the National Institutes of Health T32 training grant (T32AI134646, NIAID).

## Notes

### Competing Interest Statement

The authors have declared no competing interest.

