## Supplemental Information for "CD11c+ CD8 T cells cause IFN-γ-dependent autoimmune neuroinflammation that is restrained by PD-1 signaling"

**<u>Supplemental Figure 1</u>**

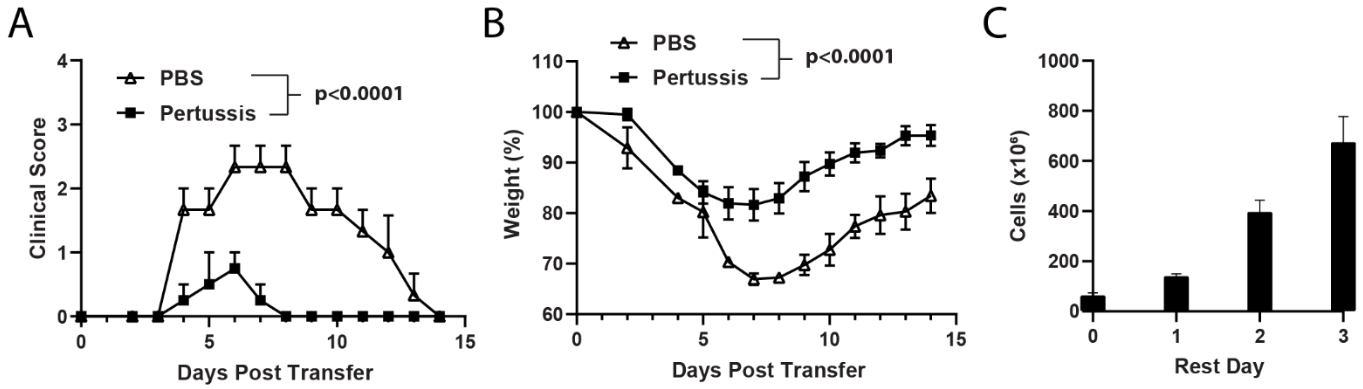


**Supplemental Figure 1. A single activation is sufficient to render Tc1 cells encephalitogenic, and pertussis toxin suppresses Tc1-mediated EAE.** Splenocytes of 8.8 mice were stimulated with $MBP_{79\text{-}87}$ for 3 days in Tc1-polarizing conditions, which were IL-12 (2 ng/mL) and IL-2 (5-10 ng/mL). After 72 h cells were rested in 10 ng/mL IL-2; each day cells were split and replated in fresh media supplemented with IL-2. On the third day, CD8 T cells were purified by MACS *via* negative selection and $2 \times 10^7$ cells in 300 µL RPMI (media only without FBS) were i.v. injected into naïve C3HeB/FeJ recipients along with 0.4 µg IL-2 per mouse. **A)** Clinical course for CD8-EAE in mice that were injected i.p. with either 200 ng pertussis toxin or PBS on day 0 and 2 post transfer (n=3 mice per group). **B)** Change in weight during CD8-EAE. Significance was determined by Two-way ANOVA. **C)** Cell number in cultures over the course of rest/expansion following 3 days activation. In the experiments shown, splenocytes from two 8.8 donor mice were used.

**Supplemental Figure 2**

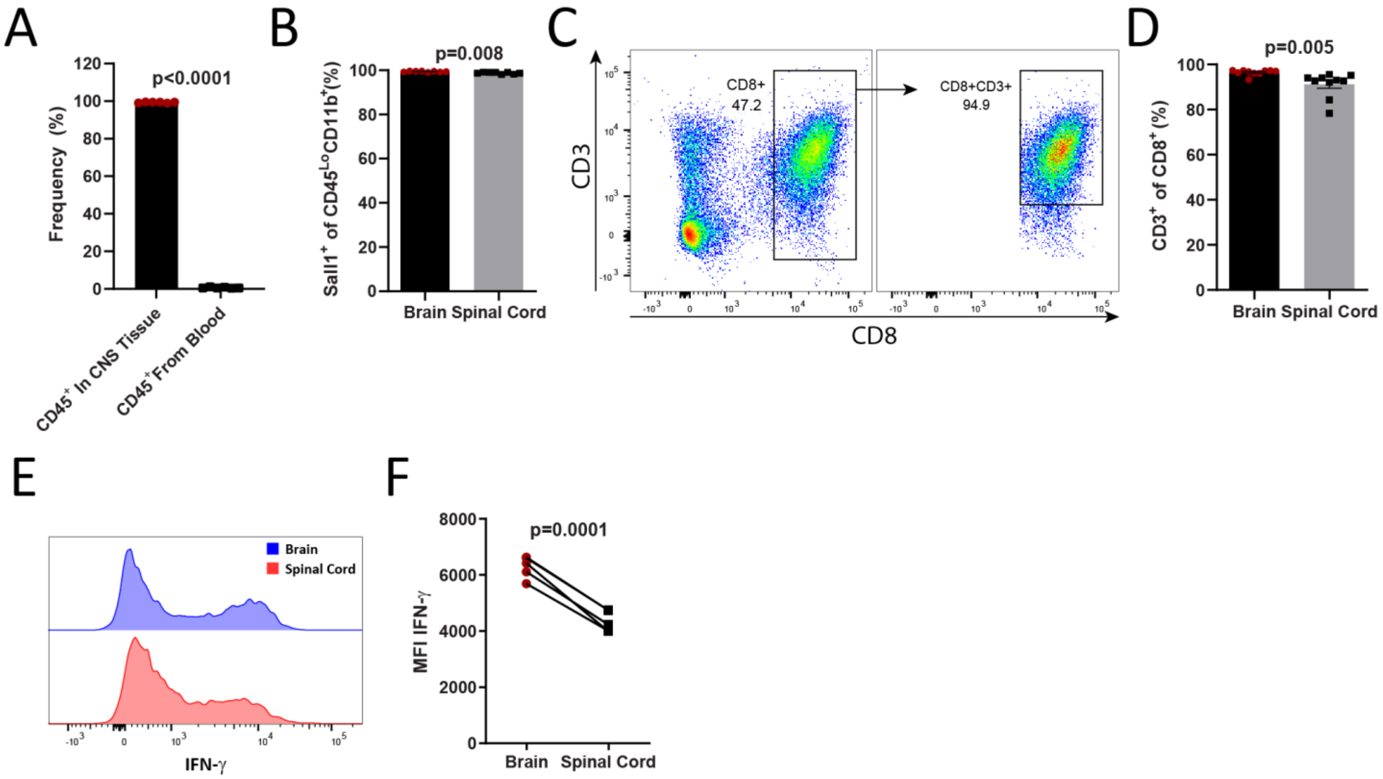


**Supplemental Figure 2. Tc1 cells cause distinct inflammation in the brain and spinal cord.** CD8-EAE was induced by i.v. transfer of 2 x $10^7$ Tc1 cells. **A)** Quantification of cells from the CNS of mice with CD8-EAE that are resident/infiltrating *vs*. those deriving from blood. On day 7 post transfer, to label CD45$^+$ cells in the blood, 10 μg of fluorochrome-labeled anti-CD45 mAb was injected i.v. 5 min prior to perfusion and CNS isolation. After isolation of cells from the CNS, cells were stained with anti-CD45 mAb conjugated to a different fluorochrome than i.v.-injected anti-CD45 mAb, and the frequency CD45$^+$ cells from CNS tissue, and CD45$^+$ cells deriving from blood were determined. **B)** Frequency of Sall1$^+$ cells among CD45$^{Lo}$CD11b$^+$ cells in the brain and spinal cord. **C)** Flow cytometry plots showing CD3 and CD8 expression among CD45$^+$Sall1$^-$ cells (non-microglial immune cells; left panel) and CD45$^+$Sall1$^-$CD8$^+$ cells (right panel). **D)** Frequency of CD3$^+$ cells among CD8$^+$ cells in the brain and spinal cord. **E)** Histograms depicting IFN-γ expression among monocytes. **F)** MFI of IFN-γ$^+$ cells in CD45$^{Hi}$CD11b$^+$Ly6G$^{Lo/-}$Ly6C$^+$ monocytes from the brain and spinal cord. Significance for A-C was determined by unpaired t test. Significance for E was determined by paired t test.

**Supplemental Figure 3**

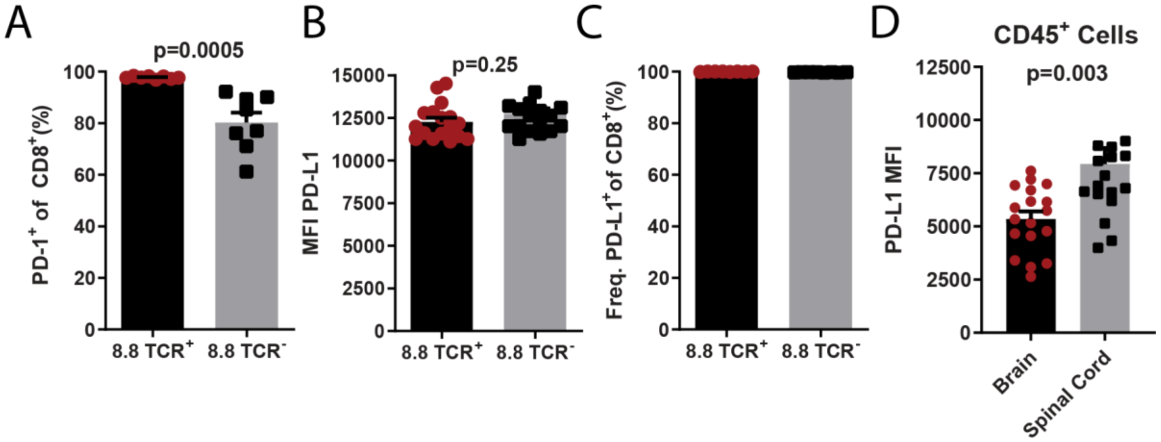


**Supplemental Figure 3. CNS CD8 T cells in CD8-EAE are predominately CD11c$^+$ with terminal effector phenotype.** CD8-EAE was induced by i.v. transfer of 2 x $10^7$ Tc1 cells and mice were sacrificed 7 days after transfer. Brain and spinal cord cells were analyzed by flow cytometry. **A)** Frequency of PD-1$^+$ cells among 8.8 TCR$^+$ and 8.8 TCR$^-$ CD8 T cells from the CNS (brain + SC). **B)** MFI of PD-L1$^+$ cells among CD8$^+$ cells. **C)** Frequency of PD-L1$^+$ cells among CD8$^+$ cells. Significance for A-C determined by unpaired t test. **D)** MFI of PD-L1 among CD45$^+$ cells. Significance was determined by paired t test.

**Supplemental Figure 4**

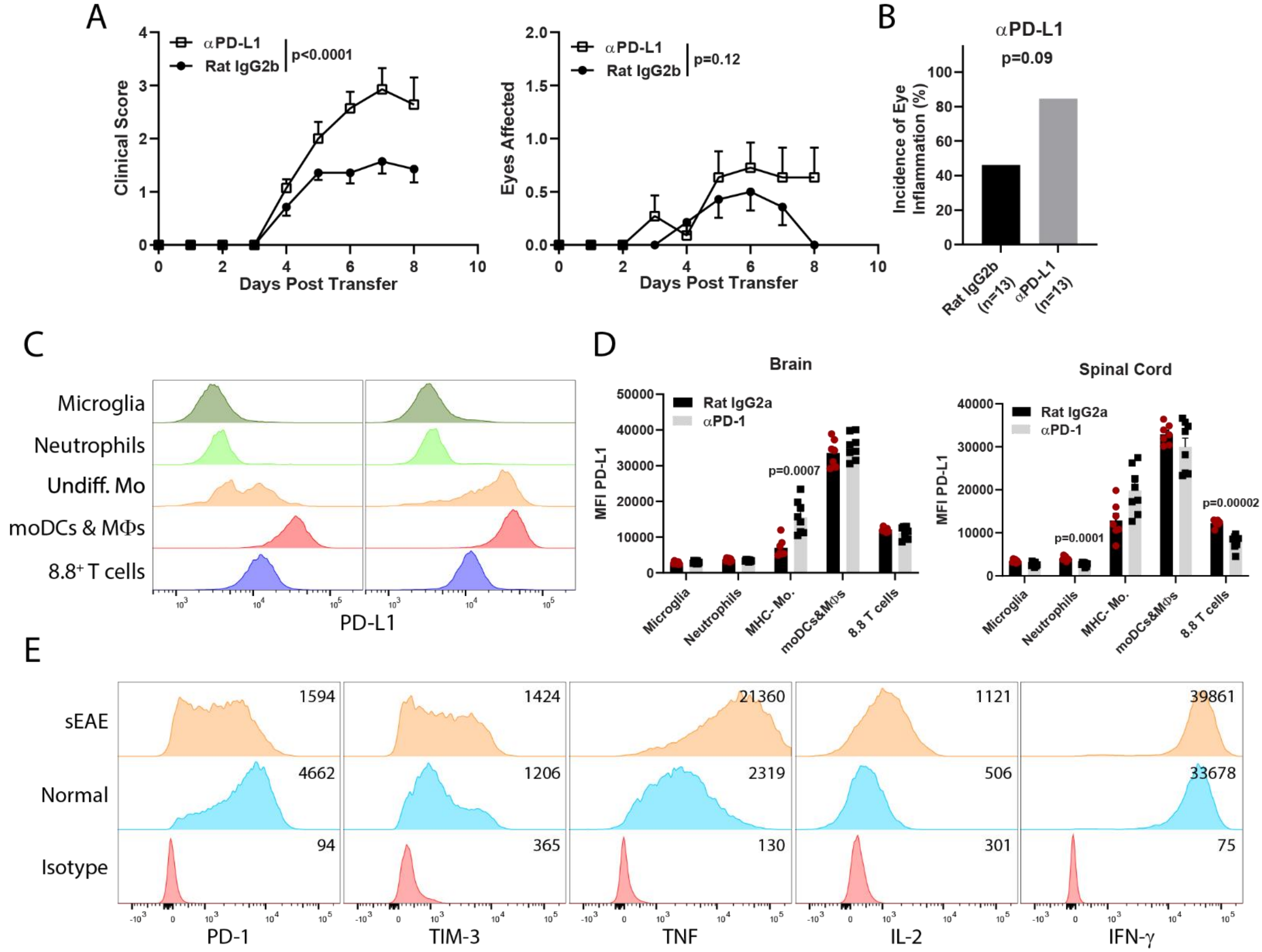


**Supplemental Figure 4. Blocking PD-L1 exacerbates CD8-EAE. A)** CD8-EAE was induced, and recipient mice were treated with 200 μg αPD-L1 or isotype mAb (Rat IgG2b) once per day starting from day of CD8-EAE induction (n=14 per group; compiled from 3 independent experiments). Statistical significance was determined by two-way ANOVA. **B)** Incidence of eye inflammation in mice treated with αPD-L1, or isotype mAb. Significance was determined by Chi-squared test. **C)** Flow cytometry histograms depicting expression of PD-1 among cell populations. **D)** MFI for PD-L1 in the brain and spinal cord among populations from F. For D statistical significance was determined by two-tailed unpaired t test. **E)** Histograms depicting expression of PD-1, TIM-3, TNF, IL-2, IFN-γ by 8.8 CD8 T cells stimulated for 3 days with $MBP_{79-87}$ that were derived from a 8.8 mouse with spontaneous EAE (sEAE) or normal appearing donors.

**Supplemental Figure 5**

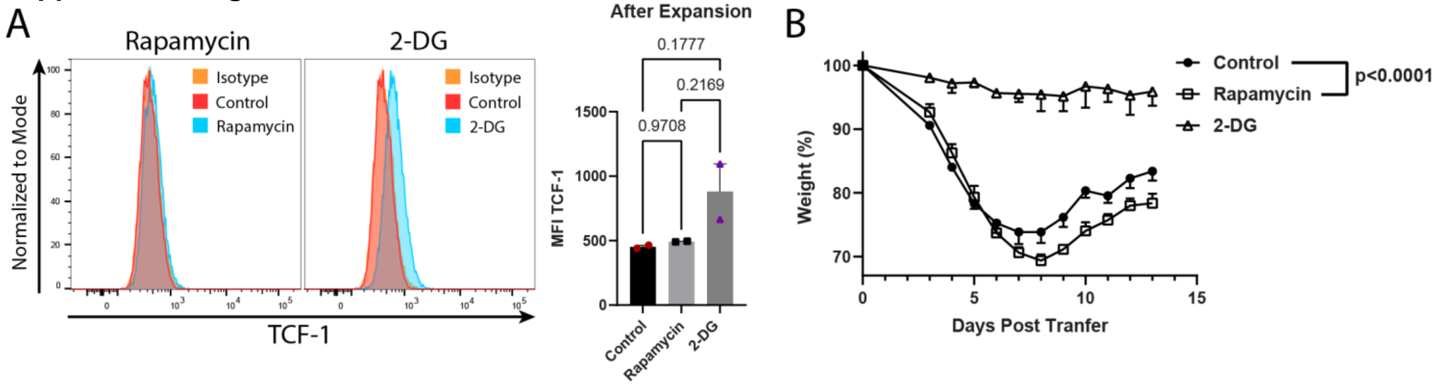


**Supplemental Figure 5. Rapamycin-treated 8.8 CD8 T cells induce greater weight loss, and 2-DG-treated cells express TCF-1. A)** Expression of TCF-1 among control, and rapamycin- and 2-DG-treated CD8 T cells after 3 day stimulation and 3 day rest. Cells were treated with 4 mM 2-DG during both activation and rest. **B)** Change in weight of CD8-EAE mice after disease was induced by transfer of control, and rapamycin- or 2-DG-treated CD8 T cells.

**Supplemental Figure 6**

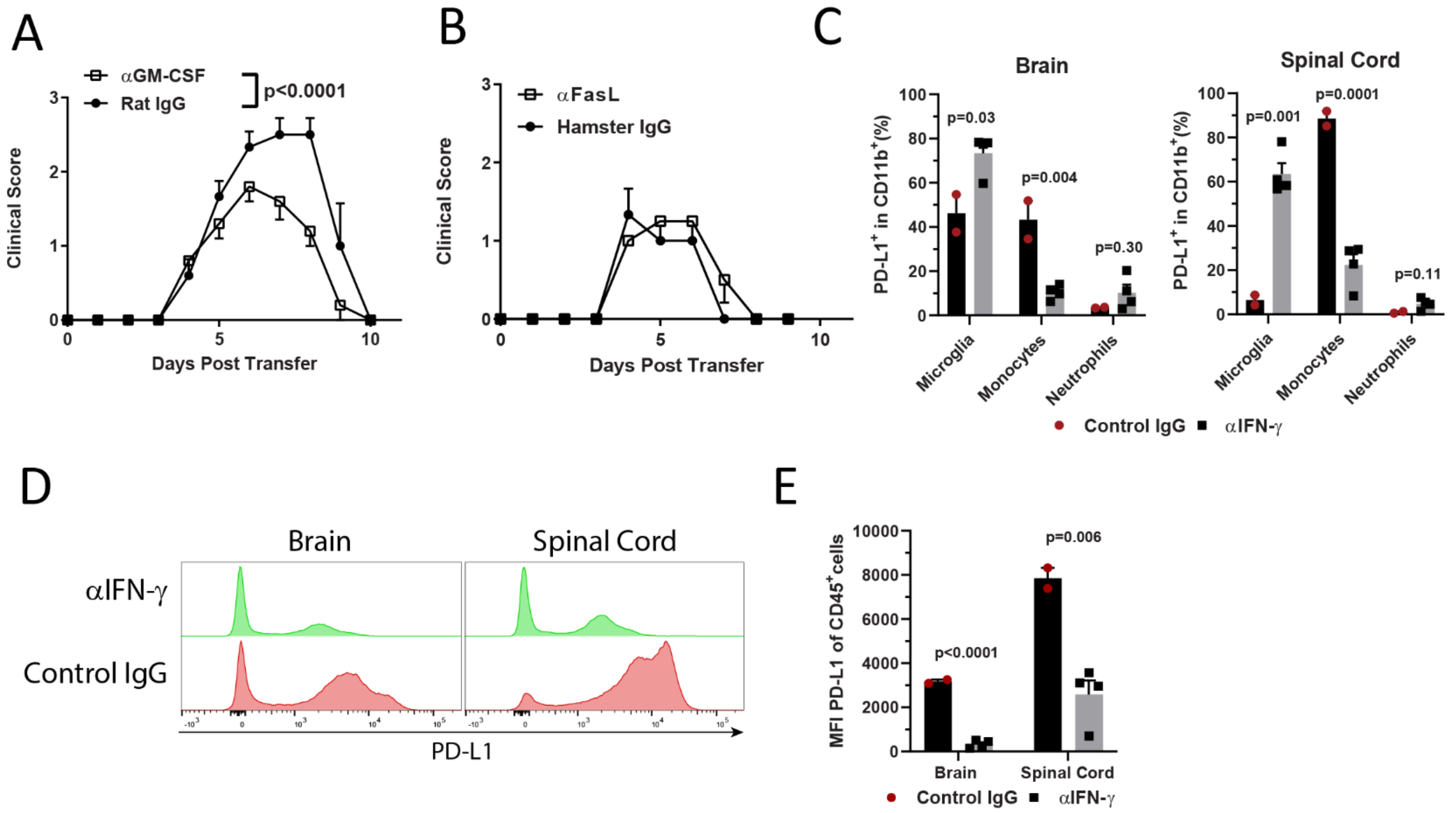


**Supplemental Figure 6. Blocking GM-CSF but not FasL suppresses EAE, and PD-L1 expression is altered by blocking IFN-γ. A)** Clinical scores of CD8-EAE mice treated with anti-GM-CSF or isotype control mAb (400 μg/day; n=5-10 per/group). **B)** Clinical scores for anti-FasL or isotype mAb-treated CD8-EAE mice (200 μg/day; n=3-4 per/group). **C)** Distribution of myeloid cells that were PD-L1$^+$CD11b$^+$ in the brain and spinal cord of anti-IFN-γ or isotype mAb-treated mice with CD8-EAE that were sacrificed on day 7 post-transfer. **D-E)** Histograms and quantification of PD-L1 MFI among CD45$^+$ cells from mice treated with αIFN-γ or control IgG.
